# Reversible in vivo regulation of drug metabolizing enzyme CYP1A2 activity through a dTAG knock-in strategy

**DOI:** 10.64898/2026.05.06.722533

**Authors:** Shenzhi Zhou, Xingyu Ji, Hanfeng Lin, Denise G. Lanza, Sung Y. Jung, Jinxia Liu, Ashish Dogra, Behnam Nabet, Kevin R. MacKenzie, Jin Wang, Martin M. Matzuk, Feng Li

## Abstract

Drug-metabolizing enzymes determine therapeutic exposure, efficacy and toxicity, but defining their isoform-specific functions in vivo remains challenging. Cytochrome P450 enzymes (P450s) are central to drug metabolism and pharmacokinetics (DMPK) and mediate the phase I metabolism of ∼75% of all marketed drugs. However, conventional knockout models can induce developmental and compensatory adaptations, and selective inhibitors are unavailable for many P450 isoforms. Here, we report the use of an inducible chemical-genetic platform for acute and specific degradation of the endogenous P450 enzyme Cyp1a2 in mice. Using CRISPR-Cas9-mediated knock-in editing, we introduced an FKBP12^F36V^ degron into the endogenous Cyp1a2 locus to generate Cyp1a2^dTAG^ mice. Treatment with the dTAG degrader dTAG-13 recruited an E3 ubiquitin ligase to CYP1A2^dTAG^, resulting in rapid and reversible proteasomal depletion of CYP1A2^dTAG^ in vivo. Temporally controlled CYP1A2^dTAG^ loss altered caffeine pharmacokinetics as expected, validating this model as a functional tool for DMPK studies. By enabling reversible suppression of drug-metabolizing enzymes without permanent deletion or chronic inhibitor exposure, this work establishes targeted protein degradation as a broadly adaptable strategy for studying drug metabolism in vivo and provides a foundation for extending inducible DMPK control to other P450s, conjugating enzymes and transporters.

## Introduction

Advancing lead compounds to clinical trials requires demonstrating preclinical efficacy and safety in animal models. Both these properties can be strongly affected by drug metabolism pharmacokinetics (DMPK), which describes the absorption, tissue distribution, metabolism, and clearance of a compound^1–3^. Despite species differences, preclinical DMPK studies in mice are widely employed and are usually broadly predictive of compound DMPK in humans^4^. Cytochrome P450 enzymes (P450s) play central roles in DMPK, in which human P450s mediate the Phase I metabolism of ∼75% of all marketed drugs^5,6^, mainly in the liver and small intestine.

Defining the isoform-specific roles of P450 enzymes is essential for understanding pharmacokinetics, drug-drug interactions, and toxicity. A drug that inhibits or enhances the activity of a P450 can cause drug-drug interactions, leading to altered efficacy or toxicity^7^. Preclinical DMPK studies of lead compounds in animals determine if a drug or its metabolites are toxic and identify the specific P450s involved, allowing adverse drug-drug interactions in patients to be predicted and avoided. Current approaches for manipulating specific P450s in vivo remain limited. Chemical inhibitors or P450-knockout (KO) animal models are the most commonly used strategies. Although selective chemical inhibitors of P450s are valuable in vitro, their application in vivo is often restricted by suboptimal potency and selectivity. Unfortunately, global knockout of a cluster of P450 genes (e.g., *Cyp3a*^-/-^) leads to compensatory changes in the expression of other P450s, with confounding effects. mRNA levels of five CYP2C family enzymes are 1.5- to 3-fold higher, and *Cyp2c55* mRNA is 30-fold higher, in the *Cyp3a*^-/-^ mouse liver compared to wild type^8^. Gene expression levels of *Cyp2a4*, *Cyp2b9*, *Cyp2b10*, and *Cyp3a13* are upregulated 6- to 48-fold in *Cyp3a*^-/-^ male mice, with smaller changes in female mice^9^. These compensatory changes in P450 expression in knock-out mouse models complicate the interpretation of studies that use these models^9^. Cyp1a2-KO rats show ∼40% lower mRNA levels of CYP2E1, CYP3A1 and liver X receptor, and CYP1A1 is upregulated ∼50-fold^10^. *Cyp1a2*^-/-^ rats exhibit a significant increase in serum cholesterol and free testosterone compared to WT, with mild liver damage, suggesting that *Cyp1a2*^-/-^ affects liver function and lipid metabolism^10^. Deleting a drug-metabolizing P450 can also affect endogenous metabolism, for example, in *Cyp3a*^-/-^ mice, liver mRNA levels of genes involved in cholesterol and bile acid metabolism (*Cyp7a1*, *Cyp8b1*, and transcription factor *Srebp-2*) are higher than in WT; hepatic cholesterol levels are 20% lower and bile acid levels are 50% higher in *Cyp3a*^-/-^ mice than in WT^11^. These changes make it difficult to interpret the data resulting from pharmacological interventions. Novel models are needed in which a given P450 can be manipulated with minimal perturbation of other P450s and of other metabolic pathways.

The dTAG (degradation tag) system enables the selective, rapid, and reversible degradation of targeted proteins in cells or organisms to probe the effects of rapid protein loss^12–16^. This chemical genetics^17^ method uses knock-in gene editing to tag a protein, sensitizing it to induced degradation by a small molecule such as a dTAG degrader. The target protein is genetically fused to the FKBP12^F36V^ degron tag, an engineered mutant form of human FK506-binding protein 12 (FKBP12). A cell-permeable dTAG molecule that binds FKBP12^F36V^ (but not FKBP12^WT^) recruits an E3 ubiquitin ligase to the tagged target protein, causing its ubiquitination and subsequent degradation by the proteasome (**Fig. 1**). The tagged protein is present until the dTAG molecule drives selective degradation. dTAG system offers target specificity, temporal control, minimal compensatory changes, reversibility and degrading targets that lack chemical probes^18–21^: FKBP12^F36V^-tagged proteins are degraded only in the presence of the dTAG molecule; subsequent metabolic clearance (washout) of the dTAG allows the tagged proteins to recover to their basal expression levels. This work uniquely adapts the dTAG system, originally developed for cancer research, to in vivo pharmacology and establishes a proof of concept for the acute and temporally controlled modulation of a key drug-metabolizing enzyme CYP1A2 (**Fig. 1**) . In this study, we report the generation and systematic characterization of *Cyp1a2*^dTAG^ mice and demonstrate their application in pharmacokinetic studies.

**Figure 1.**
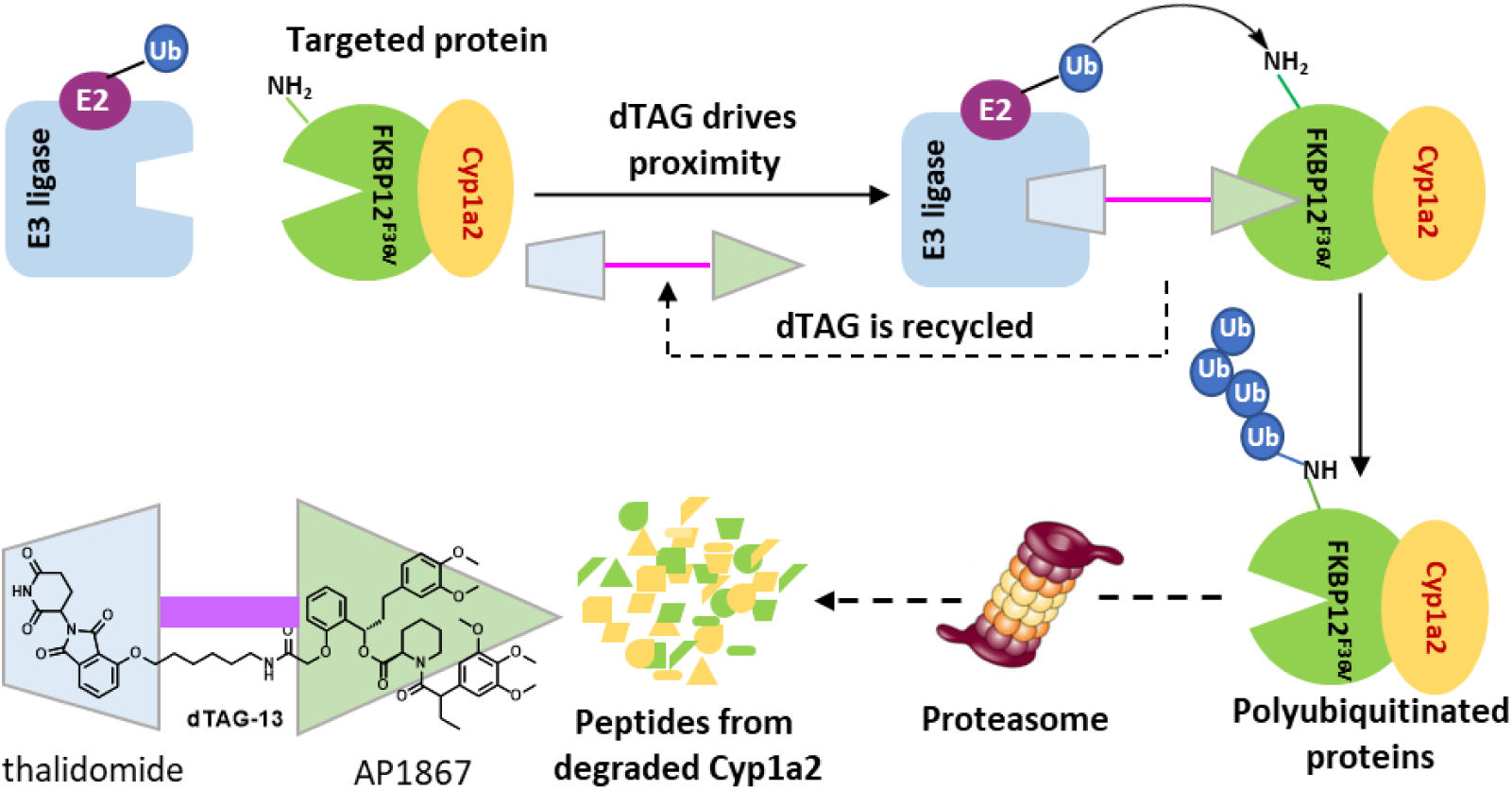
The proposed dTAG system for Cyp1a2 degradation. Bifunctional small molecule **dTAG-13** binds to the FKBP12^F36V^ domain of tagged CYP1A2 and recruits an E3 ligase; ubiquitination (curved arrow) directs fusion protein degradation. The AP1867 moiety of dTAG-13 (bottom left) binds FKBP12^F36V^ selectively; the thalidomide moiety recruits cereblon.

## Results and discussion

### Design of *Cyp1a2*^dTAG^ mouse generation

As presented in **Fig. 2**, a guide RNA (gRNA) targeting the C-terminal region of the mouse *Cyp1a2* gene was designed to introduce an FKBP12^F36V^ degron tag immediately upstream of the endogenous stop codon for C-terminal tagging of *Cyp1a2*. The selected gRNA (TGCCTCGACAATCTTCACT, PAM: TGG) targets chromosome 9 (57,677,213–57,677,235) in the C57BL/6J genome and no predicted off-target sites with fewer than three mismatches were identified within coding or intronic regions. Cleavage efficiency was verified by electroporating Cas9-gRNA complexes into one-cell-stage embryos, followed by blastocyst culture and PCR genotyping, which confirmed efficient editing at the targeted locus of C-terminal region of *Cyp1a2*. A single-stranded donor repair template was then designed for homology-directed repair (HDR). The donor template contained homology arms flanking the Cas9 cleavage site and encoded an in-frame C-terminal fusion composed of a five-amino-acid flexible linker followed by the FKBP12^F36V^ degron sequence, enabling inducible protein degradation through the dTAG system. Cas9-mediated cleavage and HDR introduced an approximately 340-bp donor sequence into the endogenous *Cyp1a2* locus. Zygotes were microinjected with Cas9 mRNA (100 ng/μL), sgRNA (20 ng/μL), and donor DNA (100 ng/μL) in RNase-free modified TE buffer to generate *Cyp1a2*^dTAG^ knock-in mice.

**Figure 2.**
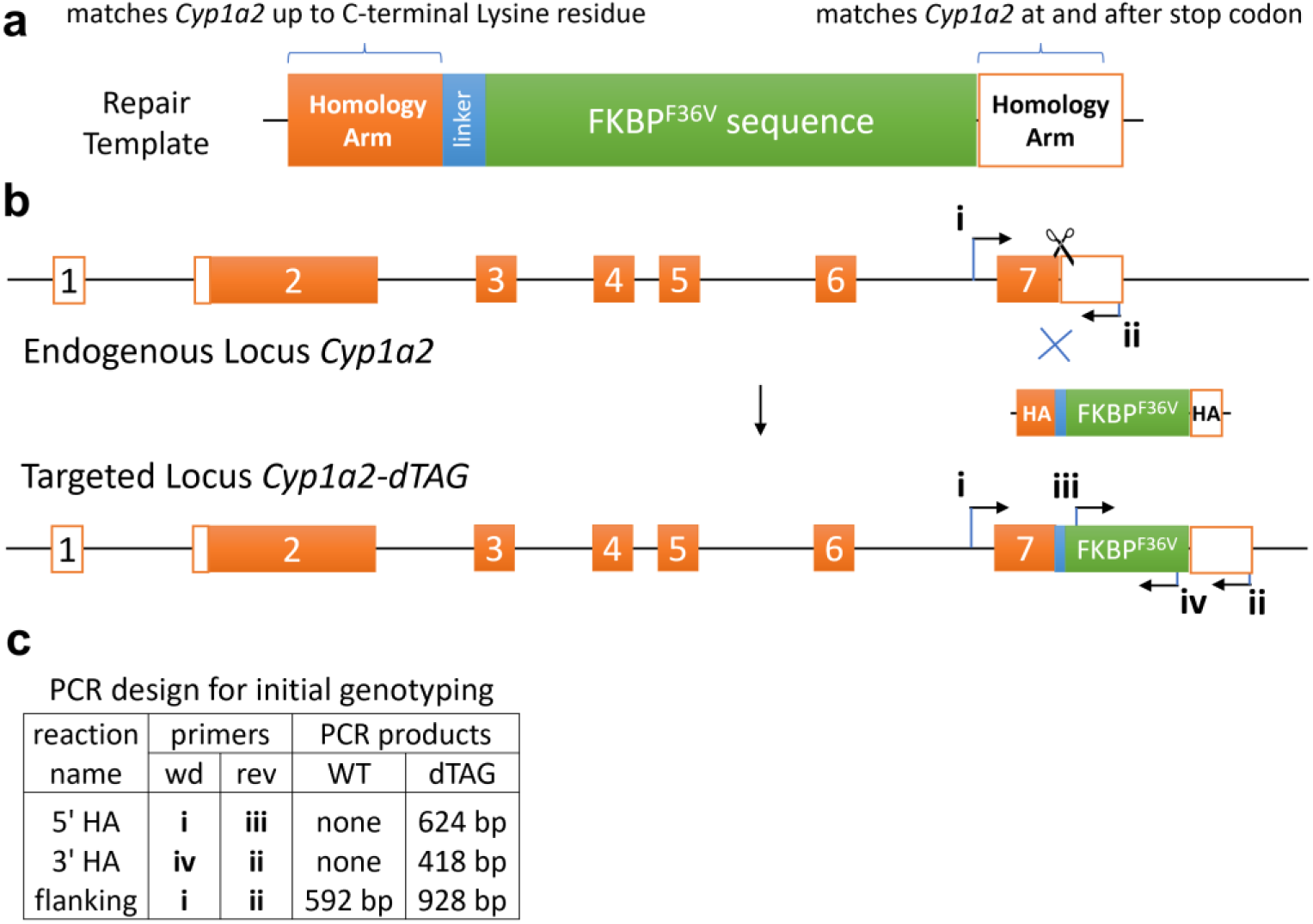
Design of dTAG knock-in at Cyp1a2 C-terminus in the germline of C57BL/6J mice. **a**, A knock-in repair template was designed to insert a five-residue linker and the FKBP12^F36V^ sequence. **b**, The *Cyp1a2* C-terminus was targeted with a gRNA proximal to the translational stop (scissors) and insertion was mediated by homologous-recombination-directed repair (blue X). Lines represent introns, orange boxes represent coding exons, white boxes represent non-coding exons. **c**, PCR-based genotyping was used to identify successfully targeted alleles with the defined primers. HA, homology arm.

The C-terminal tagging strategy was chosen to preserve endogenous *Cyp1a2* gene regulation while enabling post-translational control of CYP1A2 protein stability. Inserting the FKBP12^F36V^ degron immediately upstream of the native stop codon preserves the endogenous promoter, intronic regulatory elements, and physiological expression pattern of CYP1A2. Positioning the degron at the C terminus also avoids disruption of the N-terminal membrane-targeting sequence, which is essential for proper localization of P450 enzymes to the endoplasmic reticulum and for interactions with redox partners such as P450 reductase. To minimize steric interference, a five-amino-acid flexible linker was inserted between CYP1A2 and the degron to help maintain proper folding and catalytic function of the enzyme. In contrast, fusion of a degron to the N-terminus therefore carries a substantial risk of disrupting membrane targeting, impairing protein folding, or altering basal enzymatic activity independently of ligand-induced degradation. Because the C-terminus of CYP1A2 resides on the cytosolic side of the endoplasmic reticulum and does not mediate membrane anchoring, it is likely to be more permissive to fusion without compromising native topology. Thus, placement of the dTAG moiety at the C-terminus is expected to preserve physiological localization and catalytic competence under basal conditions while maintaining accessibility of the degron to recruit E3 ubiquitin ligases upon ligand treatment. Although C-terminal tagging may still influence protein stability or protein-protein interactions, this configuration minimizes perturbation of the critical N-terminal targeting domain and provides a more reliable platform for interpreting degradation-dependent phenotypes.

### Creation and physiological characterization of homozygous *Cyp1a2*^dTAG^ mice

The CRISPR/Cas9 assembly comprising the Cas9 mRNA, the specific sgRNA, and the DNA donor template containing the dTAG sequence, was microinjected directly into the pronuclei of fertilized mouse zygotes. These manipulated embryos were then surgically implanted into the oviducts of five pseudo pregnant foster mothers (Strain: C57BL/6J). Four of the five females were pregnant giving birth to a total of 14 pups (5 male and 9 female). They were genotyped via PCR demonstrating double strand breaks induced by Cas9 resulted in successful knock-in of the desired variant in one female individual (**Fig. S1**). Two reactions with one primer outside and one inside the donor FKBP12^F36V^ sequence were performed to verify the correct targeting of the degron allele. This mouse having both PCR products (5’ HA: 624 bp; 3’ HA: 418 bp) was identified as putative founder, which was backcrossed with stock C57BL/6NJ mice to generate F1 pups (**Fig. 3a**). All F1 mice were genotyped, and mice with positive PCR results for the two homology arm reactions as in **Fig. 2** were confirmed by Sanger sequencing. We selected C57BL/6NJ for crossing with the founder (C57BL/6J) because C57BL/6J carries a naturally occurring mutant nicotinamide nucleotide transhydrogenase (Nnt) allele, whereas C57BL/6J does not. Nnt is a mitochondrial inner membrane enzyme that supports NADPH generation and redox homeostasis through detoxification of reactive oxygen species. Because NADPH is also an essential cofactor for P450 enzymes, crossing with C57BL/6NJ may help reduce the potential impact of the C57BL/6J Nnt defect on the phenotype of interest. Evaluation of Nnt status in the *Cyp1a2*^dTAG^-HOM mice is beyond the scope of the current study; however, this will be examined in the future work.

**Figure 3.**
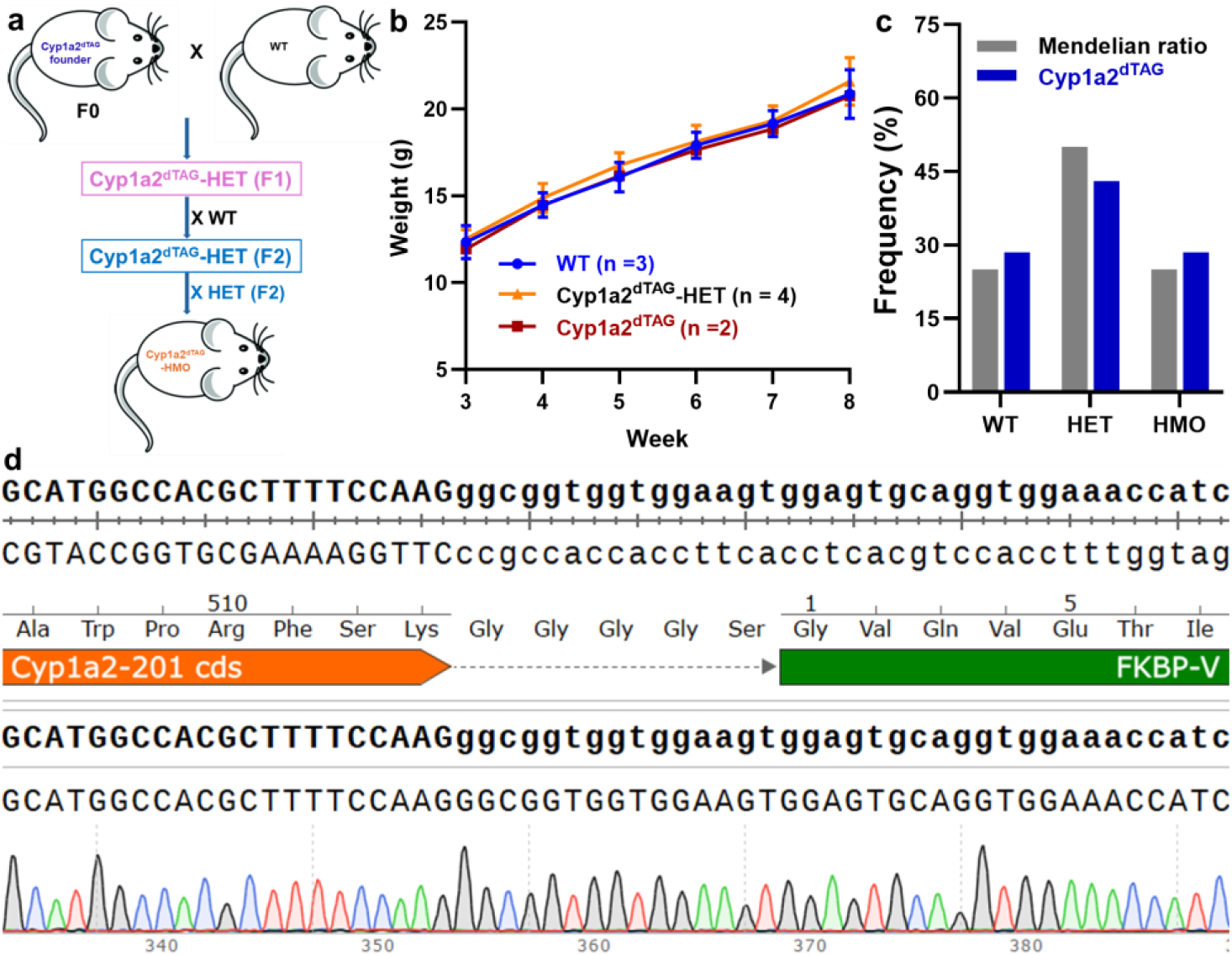
Generating Cyp1a2^dTAG^ homozygotes and physiological characterization. **a**, Flowchart for generating homozygous *Cyp1a2*^dTAG^ mice. *Cyp1a2*^dTAG^-HET, heterozygous; *Cyp1a2*^dTAG^-HMO, homozygous. **b,** The HET × HET cross gave a normal litter size and genotypes in Mendelian ratios, with no indication that homozygotes might be disfavored. **c,** *Cyp1a2*^dTAG^ does not affect growth rate. Body weight of developing WT, *Cyp1a2*^dTAG^-HET and *Cyp1a2*^dTAG^-HMO mice. The data are express as mean with standard deviations. **d,** Sanger sequencing at the site of in-frame fusion of a homozygote. A total of 66 mice were analyzed for **Figs. 3b** & **3c**.

To generate *Cyp1a2*^dTAG^ homozygous mice (*Cyp1a2*^dTAG^-HOM) and establish a stable germline-transmitted colony (**Fig. 3a**), *Cyp1a2*^dTAG^ heterozygous founder offspring (C*yp1a2*^dTAG^-HET, F1) were first crossed with WT mice to yield F2 generation *Cyp1a2*^dTAG^-HET mice and then backcrossed with F2 mice to yield F3 generation *Cyp1a2*^dTAG^-HET mice. The offspring were genotyped by standard PCR using flanking primers that distinguish the endogenous and targeted alleles (**Fig. 2c**), and correct integration at both homology arms was confirmed by Sanger sequencing. Intercrossing F3 heterozygotes generated WT (n = 17, sum from 11 cohorts), *Cyp1a2*^dTAG^-HET (n = 34, sum from 11 cohorts), and *Cyp1a2*^dTAG^-HOM (n = 15, sum from 11 cohorts) offspring. The genotype distribution from HET × HET matings followed the expected Mendelian 1:2:1 ratio (**Fig. 3c**) and representative partial sequencing data for *Cyp1a2*^dTAG^-HOM mice are shown in **Fig. 3d**. To assess postnatal development, all offspring from HET × HET crosses were weighed weekly after weaning. Body weights were monitored weekly post-weaning for a cohort of offspring. Growth curves showed that body weights of *Cyp1a2*d^TAG^-HOM mice and HET littermates did not differ significantly from those of WT, suggesting that introduction of the fusion protein did not adversely affect growth, food intake, or nutrient utilization. (**Fig. 3b**). We subsequently established a stable breeding colony using *Cyp1a2*^dTAG^-HOM mice in the F4 generation. *Cyp1a2*^dTAG^-HOM mice were viable and fertile, with no obvious abnormalities in appearance, behavior, or development observed in either heterozygous or homozygous animals throughout the observation period. In addition, sex ratios from HOM × HOM matings were generally balanced (data not shown), indicating no overt impairment in reproductive outcomes.

In addition, liver injury biomarkers alanine aminotransferase (ALT) and aspartate aminotransferase (AST) were examined, and liver morphology was evaluated by H&E staining. H&E staining of liver sections from HOM mice showed no apparent histopathological abnormalities (**Fig. S2a** and **S2b**). Plasmas ALT and AST levels were not significantly different between WT and HOM mice (**Fig. S2c** and **S2d**). Together, these findings indicate that introduction of the *Cyp1a2* fusion knock-in allele did not cause detectable liver functional or morphological changes under the tested conditions. Thus, physiologically normal homozygous *Cyp1a2*^dTAG^ mice were successfully generated and subsequently subjected to functional characterization of this model.

### Characterizations of *Cyp1a2*^dTAG^-HOM mice

We first evaluated hepatic CYP1A2 and CYP1A2^dTAG^ protein expression in WT, HET, and HOM mice by Western blot analysis (n = 4 per genotype; 2 males and 2 females) (**Fig. 4a**). As shown in **Fig. 4a**, a single band corresponding to native CYP1A2 was observed in WT mice, whereas a single higher-molecular-weight band (about 12 kDa) corresponding to CYP1A2^dTAG^ protein was observed in HOM mice. In HET mice, two distinct bands were detected, one corresponding to native CYP1A2 and the other to CYP1A2^dTAG^. The FKBP12^F36V^-tagged CYP1A2 protein (approximately 70 kDa, theoretically) was readily detected by the anti-CYP1A2 antibody and migrated above the native CYP1A2 band (approximately 58 kDa), consistent with the expected 12 kDa increase in molecular mass following FKBP12^F36V^ tagging. CYP1A2^dTAG^ protein was detected in HET and HOM mice, but not in WT mice, whereas native CYP1A2 protein was detected in WT and HET mice, but not in HOM mice, as expected.

**Figure 4.**
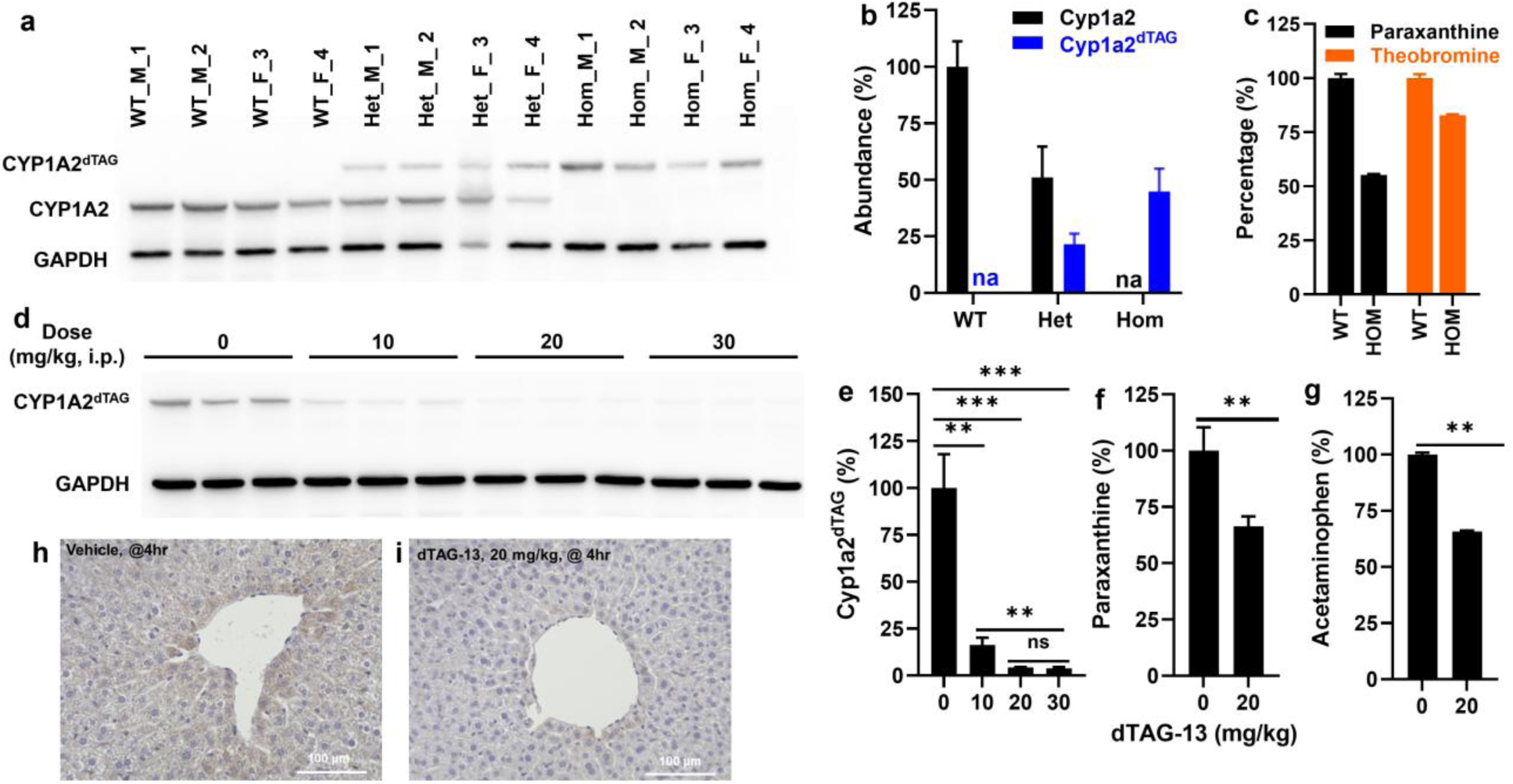
Functional characterizations of CYP1A2^dTAG^ mice. **a**, Hepatic CYP1A2 and CYP1A2^dTAG^ protein expression in WT, HET, and HOM mice. **b**, Densitometric quantification of CYP1A2 and CYP1A2^dTAG^ protein. **c**, CYP1A2^dTAG^ catalytic activity. **d**, Dose-dependent degradation of CYP1A2^dTAG^ protein mediated by dTAG-13 in HMO mice. **e**, Densitometric quantification of CYP1A2^dTAG^ protein in liver 4 hours post dTAG-13 treatment. **f** & **g**, CYP1A2^dTAG^ activity in MLMs prepared from HOM mice treated with vehicle or dTAG-13. **h**, Hepatic immunostaining of CYP1A2^dTAG^ in HOM mice (vehicle). **i**, Hepatic immunostaining of CYP1A2^dTAG^ in HOM mice (dTAG-13). For densitometric quantification in Figure 4b, native Cyp1a2 level was set as 100%; in Figure 4e, protein levels in vehicle-treated mice were set as 100%. M, male; F, Female. na, not available; ns, not significant; **, p<0.01, ***; P<0.001; HET, heterozygous; HOM, homozygous. n = 4.

Densitometric quantification showed that hepatic CYP1A2^dTAG^ protein levels in HOM mice were approximately 50% of native CYP1A2 levels in WT mice, whereas CYP1A2^dTAG^ and CYP1A2 levels in HET mice were approximately 22% and 52% of WT CYP1A2 intensity, respectively (**Fig. 4b**). This moderate decrease is not unexpected as fusion tagging may affect protein expression or stability by altering folding, processing, or degradation. Collectively, these Western blot findings confirm FKBP12^F36V^-tagged CYP1A2 protein is successfully expressed in the knock-in mice and displays the expected genotype-dependent expression pattern. CYP1A2^dTAG^ catalytic activity was then assessed using caffeine as a probe substrate in mouse liver microsomes (MLM) prepared from WT and HOM mice (**Fig. 4c**). The formation of paraxanthine, the major CYP1A2-mediated metabolite, was ∼50% of that observed in WT mice, whereas theobromine formation remained ∼85% of WT, consistent with its predominant formation mediated by CYP2E1 rather than CYP1A2. These results demonstrate that CYP1A2^dTAG^ protein retains catalytic activity, with functional activity corresponding closely to its protein abundance in the liver (**Figs. 4b** and **4c**).

Although CYP1A2 is reported to be predominantly expressed in the liver, we further examined the expression of native CYP1A2 and CYP1A2^dTAG^ by Western blot in major mouse organs from WT and *Cyp1a2*^dTAG^-HOM mice, including brain, lung, kidney, spleen, testis, and intestine. (**Figs. S3a** and **S3b**). Liver samples from WT and HOM mice were included as positive controls. CYP1A2 or CYP1A2^dTAG^ protein was detected only in the liver and not in the extrahepatic organs examined, consistent with its known liver-predominant expression pattern. To assess the degradation efficiency of the tagged fusion protein in the liver, HOM mice were treated with vehicle or dTAG-13 (a commonly used dTAG degrader) at 10, 20, or 30 mg/kg (n = 3 per group). At 4 h post-dose, 16.2%, 4.3%, and 3.8% of CYP1A2^dTAG^ protein remained after treatment with 10, 20, and 30 mg/kg, respectively, relative to vehicle-treated controls (**Figs. 4d** and **4e**). Thus, all three doses induced substantial degradation of CYP1A2^dTAG^ in the liver with 20 and 30 mg/kg producing significantly greater degradation than 10 mg/kg, whereas no significant difference was observed between 20 and 30 mg/kg. Accordingly, 20 mg/kg was used in subsequent studies. In WT mice, dTAG-13 (20 mg/kg) did not alter endogenous hepatic CYP1A2 protein levels (99.9% of vehicle control; data not shown), indicating selective degradation of the tagged fusion protein. Consistent with the reduction in protein abundance, CYP1A2 activity in MLMs prepared from *Cyp1a2*^dTAG^-HOM mice decreased by 25% and 26% after dTAG-13 treatment when caffeine and phenacetin were used as probe substrates, respectively (**Figs. 4f** and **4g**). Immunohistochemistry further confirmed efficient degradation of the tagged protein (**Figs. 4h** and **4i**) by comparing the vehicle and dTAG-13 treatment. Overall, these results demonstrate that the CYP1A2^dTAG^ system is functionally responsive to dTAG-13-mediated selective degradation in vivo, supporting successful establishment of the dTAG platform in this knock-in mouse model.

To characterize the degradation kinetics of CYP1A2^dTAG^ in vivo, HOM mice were treated with dTAG-13 at 20 mg/kg and liver samples were collected at 0, 1, 2, 4, 8, 16, 24, 32 and 48 h after dosing (n = 2 per time point). As shown in **Figs. S3c** and **S3d**, hepatic CYP1A2^dTAG^ protein levels decreased rapidly, with a reduction to 39.7% of baseline by 1 h and maximal degradation at 4 h, when only 7.3 % of the protein remained. Partial recovery was evident by 8 h, with protein levels increasing to 17.2% of baseline, and expression gradually returned to near basal levels by 24 h. Consistent with our previous report showing that dTAG-13 has a short half-life in mice (t_1/2_ = 3.1h)^22^, hepatic concentrations of dTAG-13 were very low at 8 h after dosing. Protein levels remained stable thereafter through 32 and 48 h (data not shown). These results demonstrate that dTAG-13 mediates rapid, robust, and reversible degradation of FKBP12^F36V^-tagged CYP1A2 in vivo. Because the tagged protein began to recover by 8 h after treatment, HOM mice were dosed twice with dTAG-13, at 0 and 8 h. This regimen maintained hepatic CYP1A2^dTAG^ protein levels below 10% of baseline (**Fig. S3e**), indicating that prolonged suppression is achievable over a 24-h period with two administrations. However, such a dosing strategy may be less practical for DMPK studies. Therefore, developing a degrader with a longer half-life would likely be more suitable for sustained in vivo suppression of the tagged protein in the future.

### FKBP12^F36V^ knock-in at *Cyp1a2* has minimal impact on the liver proteome

To evaluate the effect of the knock-in on the mouse liver proteome, particularly on drug-metabolizing enzymes, we next performed MS-based untargeted proteomic analysis to determine whether expression of FKBP12^F36V^-tagged CYP1A2 affects the hepatic proteome.

Proteomic analysis of liver samples collected from WT and HOM mice treated with either vehicle or dTAG-13 quantified approximately 7,000 proteins (**Fig. 5a**). Comparison of HOM and WT mice treated with vehicle showed that the global hepatic proteome remained largely unchanged, with most proteins exhibiting no substantial difference in abundance (|log2 fold change| > 1 and adjusted P < 0.01). The most significant change was reduced CYP1A2 abundance in HOM mice, remaining approximately 30% of WT level, but the fusion of FKBP12^F36V^ to CYP1A2 did not broadly perturb endogenous hepatic protein composition. Notably, the abundances of other major phase I and phase II drug-metabolizing enzymes, including CYPs, UGTs, GSTs, and transporters, were comparable between WT and HOM mice, supporting the conclusion that FKBP12^F36V^-tagged CYP1A2 has minimal impacts on other hepatic drug metabolism pathways. The lower abundance of CYP1A2^dTAG^ in HOM mice was consistent with the Western blot results.

**Figure 5.**
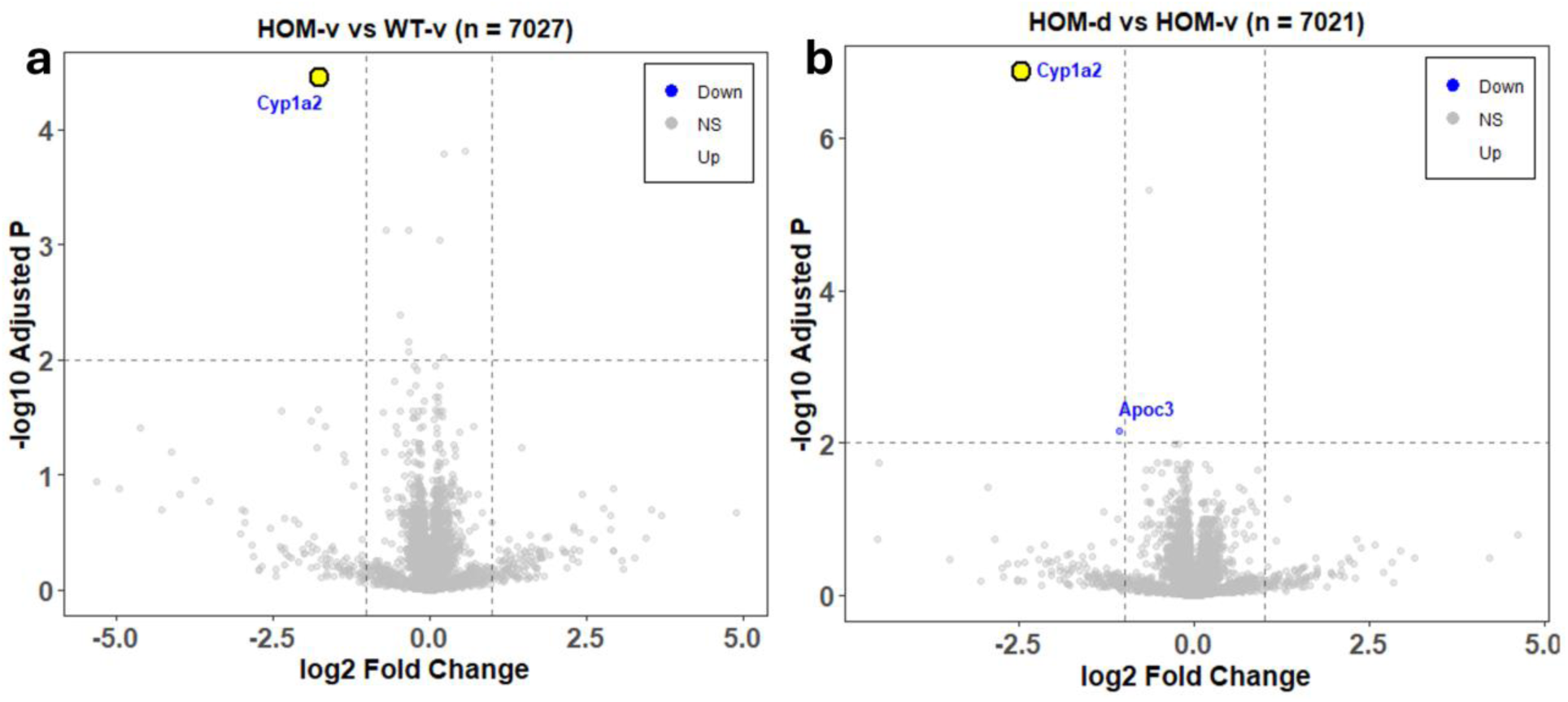
Proteomics analysis of mouse liver among WT and Cyp1a2dTAG-HOM. **a**, HOM-v vs. WT-v. **b**, HOM-d vs. HOM-v. v, vehicle; d, dTAG-13. **\***Cyp1a2 in the figure refers to CYP1A2 or CYP1A2^dTAG^. (n = 6, 3 males and 3 females).

At the transcriptional level, RT-PCR analysis of liver tissues showed that *Cyp1a2* mRNA expression was comparable between WT and HOM mice (**Fig. S4**). The mRNA levels of *Cyp3a11*, *Cyp2e1*, *Ugt1a1*, and *Abcg2* were also unaffected, indicating that insertion of the FKBP12^F36V^ tag did not alter transcriptional regulation or trigger compensatory responses. Together, the unchanged *Cyp1a2* mRNA level together with the reduced CYP1A2^dTAG^ protein level in HOM mice suggest that the lower protein level in HOM mice is primarily attributable to post-translational effects, such as altered protein stability. To our knowledge, a comprehensive baseline liver proteomic comparison between untreated *Cyp1a2*^-/-^ and WT mice has not been well established. Our proteomic analysis of liver from *Cyp1a2*^-/-^ and WT mice suggested that Cyp1a2^-/-^ significantly change the liver proteome compared to WT mice (**Fig. S5**), such as NNT (Nicotinamide Nucleotide Transhydrogenase), the most significantly downregulated protein (drops almost 65-fold), suggesting a severe disruption in mitochondrial redox homeostasis. Some muscle-related contractile proteins, including TNNC2, TNNT3, MYH4, MYL11 also decreased, while a few proteins involved in steroid and taurine metabolism were significantly upregulated by 2 fold or more, such as CSAD and AKRLC18 (|log2 fold change| >1 and adjusted P < 0.05, data not shown) indicating a chronic metabolic reprogramming of endogenous metabolism due to the loss of CYP1A2. Compared with the *Cyp1a2*^-/-^ model, the newly developed *Cyp1a2*^dTAG^ system lacks the extensive systemic reprogramming typically observed in KO model, as we had expected.

We next compared the hepatic proteomes of HOM mice treated with vehicle or dTAG-13 for 4 h (**Fig. 5b**). CYP1A2^dTAG^ protein was significantly reduced following dTAG-13 treatment, reaching approximately 18% of the level observed in vehicle-treated mice, indicating selective degradation of CYP1A2^dTAG^ in the liver. Global proteomic analysis further demonstrated the high specificity of this system, as the abundances of other drug-metabolizing enzymes, including members of the P450 superfamily, as well as proteins involved in essential hepatic functions, remained largely unchanged after dTAG-13 administration (|log2 fold change| > 1 and adjusted P < 0.01). dTAG-13 treatment also resulted in a significant decrease in APOC3, which might reflect a rapid reaction following ER-resident protein (CYP1A2) depletion, possibly trigger to prevent lipid accumulation. Overall, our results show that the *Cyp1a2*^dTAG^ model has minimal compensatory effects and that dTAG-13 can efficiently deplete CYP1A2^dTAG^ in vivo, thereby enabling acute temporal control of enzyme levels for mechanistic DMPK studies.

### Pharmacokinetic studies of caffeine in homozygous *Cyp1a2*^dTAG^ mice

Caffeine is a classic substrate of CYP1A2 and has been used to assay the metabolic activity of CYP1A2 in vitro and in vivo^10,23,24^. We first used caffeine as an in vivo probe for evaluating homozygous *Cyp1a2*^dTAG^ mice for its application in PK studies. Based on the time course of dTAG-13-mediated degradation (**Fig. S3c**), hepatic CYP1A2^dTAG^ protein levels declined by more than 40% within 1 h and remained around 50% of baseline for about 16 h. This degradation time window provided an appropriate time frame for evaluating the pharmacokinetic consequences of acute CYP1A2 loss in vivo. Accordingly, each mouse (n = 6) received a single oral dose of caffeine (10 mg/kg) 1 h after vehicle treatment and, following a 2-week washout and recovery period, received the same caffeine dose 1 h after dTAG-13 administration (20 mg/kg, i.p.). This crossover-like design allowed direct within-animal comparison of caffeine pharmacokinetics under basal and acutely depleted CYP1A2 conditions, thereby minimizing inter-animal variability.

The plasma concentration-time profiles and corresponding pharmacokinetic parameters are shown in **Figs. 6a** and **6c**. Compared with vehicle treatment, pretreatment with dTAG-13 markedly increased systemic caffeine exposure as expected (**Fig. 6a**). Specifically, *C*_max_ increased by more than twofold, from 42.80 ± 13.50 to 86.70 ± 24.99 μmol/L, and AUC increased, from 295.00 ± 63.09 to 629.20 ± 214.88 h·μmol/L. Consistent with this increase in exposure, caffeine clearance is reduced, as reflected by prolongation of the elimination half-life from 1.75 ± 0.71 to 2.84 ± 1.29 h and a decrease in apparent clearance (CL/F) from 0.04 ± 0.01 to 0.02 ± 0.00 mg/(h·μmol/L)/kg. In line with these changes, elimination half-life increased from 1.75 ± 0.71 to 2.84 ± 1.29 h, whereas apparent clearance (CL/F) decreased from 0.04 ± 0.01 to 0.02 ± 0.00 mg/(h·μmol/L)/kg. We also monitored the pharmacokinetics of the major caffeine metabolites, including paraxanthine, theobromine and theophylline (**Figs. 6b and S6a**). Among these, the exposure of paraxanthine, the principal CYP1A2-mediated metabolite, was markedly reduced after dTAG-13 treatment (**Fig. 6b**), whereas the formation of theobromine and theophylline, for which CYP1A2 plays a lesser role, was minimally affected (**Figs. S6b** and **S6c**). Overall, these findings indicate that the *Cyp1a2*^dTAG^ model enables acute depletion of CYP1A2 with minimal compensatory effects and provides a precise, pharmacologically controllable system for dissecting CYP1A2 function in drug metabolism and pharmacokinetics.

**Figure 6.**
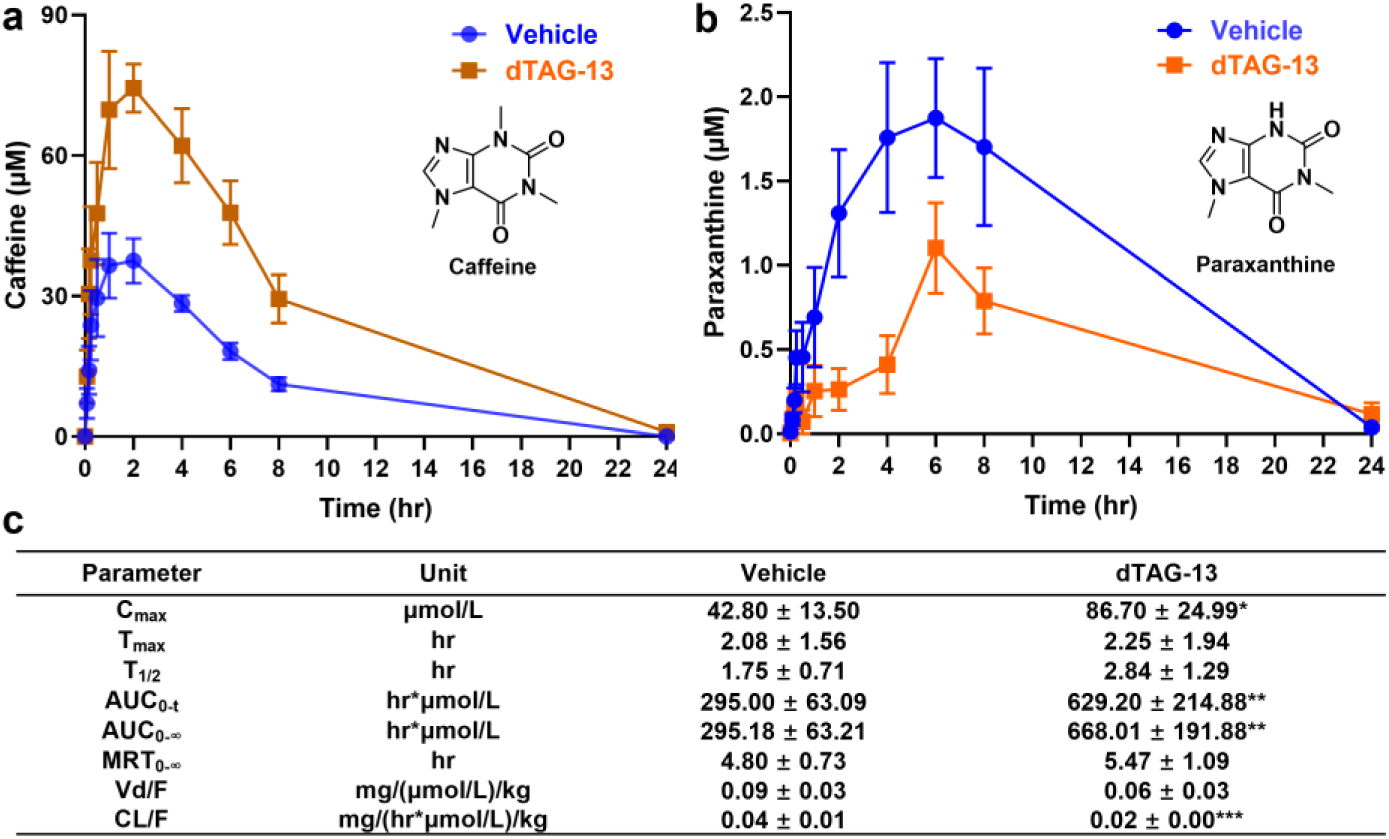
PK of caffeine in *Cyp1a2*^dTAG^-HOM mice. **a**, The PK profile of caffeine. **b**, The PK profile of paraxanthine (a CYP1A2-mediated primary metabolite). **c**, The PK parameters of caffeine. The caffeine (10 mg/kg, p.o.) was administered to mice 4 h post vehicle or dTAG-13 (20 mg/kg, i.p.). Two-week wash-out following PK of caffeine in *Cyp1a2*^dTAG^-HOM mice treated with vehicle, PK in *Cyp1a2*^dTAG^-HOM mice treated with dTAG-13 was performed. (n = 6). *, p<0.05; **, p<0.01, ***. P<0.001. HOM, homozygous.

#### Summary and future directions

Using dTAG technology to modulate the levels of drug metabolizing enzymes has the potential to transform the DMPK field by enabling rapid, selective, and reversible control of enzyme function in vivo. We have successfully generated and characterized a novel *Cyp1a2*^dTAG^ mouse model that showed limited compensatory adaptation compared with conventional *Cyp1a2*^-/-^ mice and exhibited the expected caffeine pharmacokinetic alterations following vehicle and dTAG-13 treatment, establishing proof of concept for this approach. We are currently extending this strategy to *Cyp3a11*^dTAG^ mice (**Fig. S7**), an especially important target given the central role of CYP3A in drug metabolism. In the future, we plan to further expand this platform to other major drug-metabolizing enzymes and pathways, including Cyp2d22, UGTs, and transporters, establishing a comprehensive panel of next-generation animal models for DMPK studies. We will also develop improved dTAG degraders with more favorable physiological and pharmacokinetic properties, such as longer in vivo half-life. Collectively, these efforts will create a versatile new suite of animal models for mechanistic studies of drug metabolism, pharmacokinetics, drug-drug interactions, and toxicity, thereby significantly advancing preclinical drug development.

## Experimental section

### Chemicals and materials

dTAG-13 was purchased from MedChemExpress (Monmouth Junction, NJ). Duloxetine (DLX) and agomelatine were obtained from Cayman Chemical (Ann Arbor, MI). NADPH, phenacetin, solutol, formic acid, dimethyl sulfoxide (DMSO), and RIPA lysis buffer were purchased from Sigma-Aldrich (St. Louis, MO). Water, methanol, and acetonitrile with MS grade were purchased from Thermo Fisher (Waltham, MA). Protease inhibitor cocktail (100X), ethylenediaminetetraacetic acid (EDTA), HRP conjugated secondary antibody (Cat No. 31430), ECL substrate, BCA assay kit, and 10x PBS solution were also purchased from Thermo Fisher. The 10x PBS was diluted to 1x PBS with HPLC grade water. Blotting grade nonfat milk (NFM) was obtained from Apex Bioresearch (Houston, TX). Mini PROTEAN TGX Gels (4-20%), 10x Tris/Glycine/SDS buffer, Precision plus protein™ all blue standards, and trans-blot turbo RTA mini 0.2 µm nitrocellulose transfer kit was obtained from Bio Rad (Hercules, CA). Anti-CYP1A2 primary antibody (Cat No. ab22717) was purchased from Abcam (Boston, MA). Anti-GAPDH primary antibody (Cat No. HRP-60004) was purchased from Proteintech (Rosemont, IL).

### Generation of Cyp1a2^dTAG^ knock in mouse model

*Cyp1a2*^dTAG^ mouse model (C57B/6J background) was constructed via the CRISPR-Cas9 technology by the Baylor College of Medicine Genetically Engineered Rodent Models Core. Briefly, to generate the *Cyp1a2*^dTAG^ knock-in model, a CRISPR/Cas9-mediated homology-directed repair (HDR) strategy was employed to fuse an FKBP12^F36V^ (dTAG) reporter to the C-terminus of the endogenous *Cyp1a2* gene. A specific sgRNA was designed to target the stop codon region of the *Cyp1a2* gene. The donor template was constructed to include a glycine-serine (GS) rich flexible inserted between the last codon of CYP1A2 (Phe-Ser-Lys) and the starting sequence of FKBP12^F36V^ (Gly-Val-Gln), ensuring proper protein folding and accessibility of the dTAG ligand. The donor was flanked by a 5’ homology arm and a 3’ homology arm. C57BL/6J female mice (24-32 days old) were superovulated with 5 IU PMSG followed by 5 IU hCG 48 h later, and subsequently mated with C57BL/6J males. Fertilized zygotes were collected at 0.5 dpc. The RNP complex and donor DNA were delivered into the pronuclear stage zygotes via electroporation using a BioRad Gene Pulser with the following 30 V, 12 × 1 ms pulses. Electroporated embryos were then surgically transferred into the oviducts of pseudopregnant ICR female mice. Potential founders (F0) were initially screened by PCR using a three-tier primer system. Successful 5’ junctional integration was confirmed by a genomic forward primer (5’-CACAAAAGGAACACAAAGGAAAGG) and a dTAG-specific reverse primer (5’-TCAGTTTGGCTCTCTGACCC), yielding a 624 bp fragment. The 3’ junction was verified using a dTAG forward primer (5’-ATCCTCCCGGGACAGAAAC) and a genomic reverse primer (5’-CAGGGTAGGAGGAATCTTAAATCA), yielding a 418 bp fragment. Internal integration of the CYP1A2-FKBP12^F36V^(CYP1A2^dTAG^) fusion was confirmed by a 253 bp amplicon. The precision of the integration and the preservation of the open reading frame across the junction were further validated by Sanger sequencing, confirming the seamless transition from the *Cyp1a2* C-terminus through the GS-linker to the dTAG sequence. The putative F0 founder was backcrossed with wild-type C57BL/6NJ mice to generate F1 offspring. *Cyp1a2*^dTAG^-HET mice were sequence-confirmed by the BCM GERM Core and then successively backcrossed with wild-type C57BL/6NJ mice for two generations to obtain heterozygous F3 offsprings. The heterozygous F3 mice were then intercrossed to produce homozygous *Cyp1a2*^dTAG^-HOM mice (F4). Colony maintenance genotyping was performed by Transnetyx Biotechnology (Cordova, TN). A stable colony was subsequently established from the homozygous F4 mice, and offsprings from this colony were used for the following experiments.

### Animal Treatment

Animals were housed under standard conditions with a 12-hour light/dark cycle and provided ad libitum access to food and water. All experimental procedures were conducted in strict adherence to animal study protocols approved by the Baylor College of Medicine Institutional Animal Care and Use Committee. All mice used for drug administration were between 8 and 15 weeks of age.

#### Physiological characterization

The WT, HET and HOM mice were generated from HET x HET crossing were weighed weekly after weaning to monitor their growth to assess whether their body weight of HOM mice differed from those of WT and HET mice. Their genotype distributions were also examined to determine whether genotype distributions were generally balanced and meet the Mendelian inheritance. Their appearance including fur, activity, abnormalities, and impairments were not observed in heterozygous and homozygous *Cyp1a2*^dTAG^ mice.

#### Formulations

dTAG-13 (20.0 mg/mL in DMSO) was dissolved in the formulation of DMSO/Solutol/H_2_O (10:10:80, v/v, referred to as vehicle) to yield a concentration of 2.0 mg/mL for mouse treatment. Caffeine was directly dissolved in H_2_O to a final concentration of 1.0 mg/mL for mouse treatment. All the dose solutions were prepared freshly before the treatment to avoid possible self-degradation of compounds.

#### Evaluation of liver toxicity

As CYP1A2 is mainly expressed in liver, the possibility that tagged CYP1A2 causes unintended impairment of liver was assessed by measuring plasma ALT and AST levels among WT and *Cyp1a2*^dTAG^ HOM mice. ALT and AST were measured using Cayman kits (Cayman Chemical, No. 700260 & 701640). In addition, liver morphology was examined by H&E staining. Freshly collected liver tissues were first fixed in 4% neutral buffered formalin for 24 hrs. After fixation, the tissue is dehydrated through a graded series of ethanol (from 70% to 100%) to remove water, then cleared in xylene to make it miscible with paraffin. The cleared tissue is then infiltrated with molten paraffin wax, at about 60 °C, and finally embedded in a paraffin block. Thin sections about 4 µm were cut for staining and microscopic analysis.

#### Tissue expression of CYP1A2^dTAG^ protein

The tissue including liver brain, lung, spleen, kidney, intestine, testis and epididymis from WT, HET, and HOM mice (12–15 weeks old) treated with vehicle were collected (n = 4, two males and two females) anesthetized with isoflurane. For HOM mice, an additional group treated with dTAG-13 (20 mg/kg, i.p.) were used for tissue collection. Four hours after administration, the mice were anesthetized with isoflurane, and blood was collected by cardiac puncture prior to euthanasia. All the harvested tissues were first rinsed with 1X PBS, and processed for western blotting according to the protocol of subsequent analyses. For the liver tissues, a portion of liver were fixed in 4% neutral-buffered formalin at room temperature for immunohistochemical staining and the rest of liver tissues were rapidly frozen in liquid nitrogen and stored at -80 °C for proteomics, and PCR analyses.

#### Western blot

The expression of liver CYP1A2^dTAG^ protein was first assessed using western blots. Liver samples were cut into pieces (around 30 mg-50 mg) and added to 400 µL of RIPA lysis buffer supplemented with protease inhibitor cocktail with EDTA and then homogenized manually on ice using 1.5 mL or 2 mL pestles (Axygen, Hayward, CA). Homogenized lysates were further incubated on ice for 45 min and then centrifugated at 15,000 x rcf at 4 °C for 20 min. The supernatants of each sample were collected and total protein concentration was determined at 1:50 dilution by the BCA method using a CLARIOStar Plus Microplate Reader. Thirty μg of protein for each sample were separated by SDS PAGE using 4-20% Mini PROTEAN TGX Gels. Separated proteins were transferred to a nitrocellulose membrane using the BioRad Trans-Blot Turbo and Trans-Blot Turbo RTA Transfer Kit. Membranes were blocked with 5% NFM dissolved in TBST (20 mM Tris, 150 mM NaCl, 0.1% Tween 20, pH 7.6) for 1 hr at room temperature with gentle shaking. Blocked membranes were probed for CYP1A2 (1:1000 in 5% NFM), and GAPDH (1:2500 in 5% NFM) over-night at 4 °C with gentle shaking. Membranes were washed with TBST for 8 min for a total of three washes then incubated with HRP-conjugated secondary antibodies (1:2000 in NFM) for 1 hr at room temperature with gentle rocking. Membranes were washed with TBST for 8 min for a total of three washes and then imaged by chemiluminescence using the Pierce ECL Western Blotting Substrate (Thermo Fisher Scientiffc) and a Bio Rad ChemiDoc MP Imaging System. To determine whether organs other than the liver express CYP1A2, we examined CYP1A2 protein levels in the brain, lung, spleen, kidney, intestine, testis, and epididymis. Tissue samples from these organs were processed and analyzed by western blot following the same procedures described above.

### Immunohistochemical analysis of Cyp1a2^dTAG^ in HOM mice

CYP1A2^dTAG^ protein levels and distribution were also evaluated by immunohistochemical analysis. Liver tissues were fixed overnight in 4% phosphate-buffered formaldehyde, followed by dehydration and paraffin embedding. Serial sections (4 μm) were prepared for immunohistochemical analysis of CYP1A2. Paraffin-embedded sections were first baked at 65°C for 30 min, followed by deparaffinization in three changes of xylene (5 min each) and rehydration through a graded ethanol series (100% and 95%, 10 min each). After washing in distilled water, slides were boiled in 10 mM citrate buffer (pH 6.0) for 10 min and washed. Then the sections were incubated in 0.3% hydrogen peroxide for 30 min at room temperature. Sections were blocked for 1 hr and then incubated overnight at 4°C with the primary antibody against CYP1A2 (Abcam, ab22717) diluted in blocking buffer. Following three washes in TBST, sections were incubated with a biotinylated secondary antibody for 30 min, followed by the VECTASTAIN Elite ABC Reagent (Vector Laboratories, Newark, CA) for an additional 30 min. Immunoreactive signals were visualized using a DAB substrate kit. Finally, sections were stained with Hematoxylin QS (Vector Laboratories), dehydrated through graded alcohols and xylene, and mounted and then imaged.

#### Dose-dependence and time-course for CYP1A2^dTAG^ protein degradation profiles

To determine whether the dTAG platform was successfully established, two groups of HOM mice (n = 4, two male and two female mice aged 12–15 weeks) were treated with vehicle or dTAG-13 (a dTAG degrader, 20 mg/kg, i.p.), respectively. Four hours after treatment, all the mice were anesthetized and sacrificed, and liver samples were collected and processed as described above. To establish the dose dependence, one set of HOM mice was treated at the dose of dTAG-13 (10, 20, and 30 mg/kg, i.p.; male, n = 3), and livers were collected 4 hours post-administration. An additional set of HOM mice was treated with dTAG-13 (20 mg/kg, i.p.; male, n = 2 per time point). The mice were anesthetized and sacrificed at different time points (0, 1, 2, 8, 16, 24, 32, and 48 hours) for liver sample collection.

#### CYP1A2^dTAG^ activity assay

The function of liver CYP1A2^dTAG^ protein enzyme activity was also evaluated by MLM incubation. Liver microsomes were prepared via differential centrifugation using freshly collected liver of mice given vehicle solutions or dTAG-13 (20 mg/kg) for 4 h. Briefly, mouse liver tissues (approximately 200 mg) were homogenized in ice-cold Buffer A (0.1 M phosphate buffer, pH 7.5, containing 0.25 M sucrose and 0.154 M KCl) supplemented with protease inhibitors. All procedures were performed at 4°C. The homogenate was first centrifuged at 10,000 × g for 25 minutes. The resulting supernatant was further centrifuged at 105,000 × g for 60 minutes to pellet the microsomal fraction. After washing with Buffer B (0.1 M phosphate buffer containing 20% glycerol), the microsomal pellet was resuspended in the same buffer using a manual homogenizer and then diluted after protein concentration measurement and finally stored for subsequent analysis. Several groups of MLM from different origins were employed for incubation, including WT mice or *Cyp1a2*^dTAG^-HOM mice treated with vehicle solution or dTAG-13 (20 mg/kg) for 4 h. Incubations were conducted in 1× PBS, (pH 7.4) containing 10 μM of caffeine or 20 μM of phenacetine, MLM (final concentration: 1 mg protein/mL) and then NADPH (final concentration: 1.0 mM) was added to initiate the reactions. After 15 min incubation at 37 °C with gentle shaking, reactions were terminated by addition of ice-cold acetonitrile. The mixtures were then centrifuged at 15,000 rcf for 15 min. The supernatants were collected for LC-MS/MS analysis.

#### Proteomic studies of mouse liver

About 30 to 50 mg frozen liver tissues were homogenized manually on ice in 10 volumes (w/v) of lysis buffer containing 200 mM HEPES, 0.25% SDS, and 1x Protease Inhibitor. After incubation for 45 min on ice, the lysates were centrifuged at 20,000×g for 15 minutes at 4 °C. The supernatant was collected and protein concentrations were determined by the BCA assay. Reduction was performed by adding 5 µL of 50 mM DTT and incubating at 37 °C for 15 min, and alkylation was achieved by adding 14 µL 100 mM iodoacetamide and incubating in the dark for 30 min at room temperature. The protein samples were cleaned up using SP3 method. Briefly, 30 µg protein samples and 300 µg SP3 beads (1:1 mixture of Sera-Mag Carboxylate SpeedBeads Cat. #45152105050250 and Cat. #65152105050250) were mixed. An equal volume of 100% EtOH was added to achieve 50% v/v EtOH to initiate protein binding with vigorous vortex and quick bath sonication. The mixture was incubated with 1,000 rpm shaking for 10 min at room temperature. Supernatant was removed and the beads were washed 3 times with 80% EtOH. The beads were resuspended in 50 µL of 200 mM HEPES (pH 8.5) containing 0.6 µg trypsin/Lys-C (Promega, V5073) and incubated at 37 °C with 1,100 rpm shaking overnight. The samples were then desalted and purified using StageTips (CDS Analytical, 6091). The tips were first activated with 80% acetonitrile (ACN)/0.1% formic acid and equilibrated with 0.1% formic acid. The acidified peptide digests were loaded onto the tips, which were subsequently washed twice with 0.1% formic acid. Peptides were eluted using 100 µL of 80% ACN/0.1% formic acid. The eluted peptides were dried in a speed vacuum concentrator and subsequently resuspended in 30 µL of 5% ACN/0.1% formic acid. The resulting peptides were acidified with 1 μL 100% formic acid, and 200 ng was loaded in trap and elute mode using the Thermo Scientific™ Vanquish™ Neo UHPLC system.

#### Pharmacokinetics of caffeine in Cyp1a2^dTAG^-HOM mice

The caffeine pharmacokinetics studies were conducted using Cyp1a2^dTAG^-HOM mice. For PK study, *Cyp1a2*^dTAG^-HOM mice (n = 6, 3 male and 3 female, 8-12 week old), were pre-treated with vehicle (DMSO/Solutol/H_2_O = 10:10:70, v/v; i.p. injection); after 1 hour, then orally treated with caffeine (1.0 mg/ml in water) at the dose of 10 mg/kg. About 20 μL of blood was collected via the tail vein into sodium heparin vials at 0, 0.083, 0.167, 0.25, 0.5, 1, 2, 4, 6, 8, and 24 hours post dose. The blood samples were centrifuged at 2,000 rcf for 2 min at 4 °C to separate the plasma. The samples were stored at -80 °C before analysis. After a 2-week recovery period, the mice were pretreated with dTAG-13 (2.0 mg/ml DMSO/Solutol/H_2_O = 10:10:70, v/v, i.p.) at the dose of 20 mg/kg; and after 1 hour, the mice were treated with caffeine (dissolved in H_2_O, 10 mg/kg, p.o.) and blood samples were collected, stored as above.

### UHPLC-MS analysis of PK samples

The plasma samples for LC-MS analysis were prepared following our previously established protocols ^25^. In brief, plasma samples (5 μL) were added with 30 μL of ice-cold methanol. After vortexing, the mixtures were centrifuged at 15,000 rcf for 15 minutes. Finally, a 5 μL aliquot of each supernatant was analyzed using ultra-high-performance liquid chromatography (UHPLC) coupled with a TSQ Quantis™ Triple Quadrupole mass spectrometer (Thermo Fisher Scientific, San Jose, CA). The system was equipped with a HSS T3 C-18 column (50 mm × 2.1 mm, 1.7 µm; Waters, Milford, MA) and the column temperature was maintained at 40 °C. The flow rate was set to 0.3 mL/min with a gradient elution ranging from 2% to 95% aqueous acetonitrile containing 0.1% formic acid in a 5-minute run. Ultra-pure nitrogen was employed as the sheath gas (50 arbitrary units), aux gas (10 arbitrary units), sweep gas (1.0 arbitrary unit), and collision gas. The capillary temperature was set to 325 °C, and the capillary voltage was adjusted to 3.7 kV. MS data were acquired using electrospray ionization in positive mode over a precursor/product *m/z* of 195.09 Da/110.012 for caffeine, 181.125/111.071 for paraxanthine, 181.125/96.125 for theophylline, and 244.22/170.125 for internal standard agomelatine. The chromatograms and mass spectra were acquired by Xcalibur software (Thermo Fisher Scientific, San Jose, CA). The plasma concentrations of caffeine, paraxanthine, and theophylline were calculated according to their corresponding standard curves.

### UHPLC-MS analysis of proteomics samples

For proteomics, 200 ng of resulting peptides from each sample condition was loaded in trap and elute mode using the Thermo Scientific™ PepMap™ Neo Trap Cartridge (P/N 174500) onto an EASY-Spray 150 μm * 15 cm UHPLC column (Thermo Scientific, ES906) with the Thermo Scientific™ Vanquish™ Neo UHPLC system. The sample loading, trap column washing (800 bar), and separation column equilibration (1500 bar) were performed using the Vanquish Neo UHPLC system. Buffers A and B consisted of 0.1% formic acid in water (Fisher Scientific, Optima LC-MS grade) and 0.1% formic acid/80% acetonitrile (Fisher Scientific, Optima LC-MS grade), respectively. The column was heated to 60 °C. Peptides were eluted using a 6-minute gradient from 4-35% buffer B with a flowrate of 2.5 μL/min. The eluted peptides were ionized using a Thermo Scientific™ EASY-Spray™ Source and analyzed on an Orbitrap Astral MS using narrow window DIA analysis. Briefly, MS1 spectra were collected in the Orbitrap mass analyzer every 0.6 seconds at a resolving power of 240,000 at *m/z* 200 over an *m/z* range of 380–980. The MS1 normalized automated gain control (AGC) target was set to 500% (5e^6^ charges) with a maximum injection time of 3 ms. Precursor ions were fragmented using high-energy collisional dissociation (HCD) with 25% normalized collision energy. Fragment ions were scanned in data-independent acquisition (DIA) mode in the Astral mass analyzer over an *m/z* range of 380–980 with a 2 *m/z* isolation window. The normalized AGC target was set to 500% and the maximum injection time was set to 5 ms. Window placement optimization was enabled.

#### Data analysis

For pharmacokinetic study, PK parameters of caffeine and its metabolites including half-time (T_1/2_), area under the plasma concentration-time curve during the period of observation (AUC_0–t_), area under the plasma concentration–time curve from zero to infinity (AUC_0–∞_), clearance normalized by bioavailability (CL/F), apparent volume of distribution normalized by bioavailability (V_d_/F), and the mean residence time from zero to infinity (MRT_0–∞_) were calculated in PK Solver by noncompartmental analysis. The plasma concentration–time curves were plotted in Prism 10 (GraphPad, San Diego, CA) as mean ± S.E.M. Two-tailed Student’s t-test was used to calculate the significance of difference between vehicle and dTAG-13 treated Cyp1a2^dTAG^-HOM groups. P-values less than 0.05 were considered to be statistically different (*p<0.05, **p<0.01, ***p<0.001).

For proteomics data analysis, the raw data from mass spectrometer was processed using DIA-NN software (v1.9.2 on Linux), searched against the Mus musculus proteome database downloaded from UniProt (March 2025). Both peptide-spectrum matches (PSMs) and proteins were filtered at a 1% false discovery rate (FDR). The resulting protein intensities matrix was exported and plotted using in-house python script which performs limma statistics to correct for multiple test FDR (shown as adjusted P).

## Abbreviations

P450: cytochrome P450
dTAG: degradable tag
FKBP12^F36V^: phenylalanine 36 to valine mutant of the 12-kDa FK506-binding protein
DMPK: drug metabolism and pharmacokinetics

