## Supplemental figures for "Reversible in vivo regulation of drug metabolizing enzyme CYP1A2 activity through a dTAG knock-in strategy"

‖, contribute equally to this work.

**Corresponding Author**

Feng Li – Center for Drug Discovery, Baylor College of Medicine

One Baylor Plaza, Houston TX, USA

orcid.org/0000-0002-9680-4614

**Figure S1.** Identifying a putative founder using PCR-based genotyping

**Figure S2.** Evaluation of the effects of Cyp1a2$^{dTAG}$ on liver using H&E staining and ALT&AST assay.

**Figure S3.** The expression of CYP1A2 and CYP1A2$^{dTAG}$ in major mouse organs and turnover rate of CYP1A2$^{dTAG}$ protein

**Figure S4.** Relative abundance of mRNA in WT and *Cyp1a2*$^{dTAG}$ -HOM mice

**Figure S5.** Proteomics analysis of mouse liver among WT and *Cyp1a2*$^{-/-}$ mice.

**Figure S6.** PK of theobromine and theophylline (CYP1A2-involved caffeine metabolites) in *Cyp1a2*$^{dTAG}$ -HOM mice

**Figure S7.** Design of Cyp3a11 dTAG knock-in.

**Figure S1. Identifying a putative founder using PCR-based genotyping.**

| Mouse ID | Gender | 5'Homology Arm | 3'Homology arm | Internal | Genotyping Result |
|---|---|---|---|---|---|
| 4294-001 | MALE | | | no band | |
| 4294-002 | MALE | | | band | |
| 4294-003 | MALE | | | no band | |
| 4294-004 | MALE | | | band | |
| 4294-005 | MALE | | | band | |
| 4294-006 | FEMALE | band | band | two bands | putative founder |
| 4294-007 | FEMALE | | | band | |
| 4294-008 | FEMALE | | | no band | |
| 4294-009 | FEMALE | | | no band | |
| 4294-010 | FEMALE | | | no band | |
| 4294-011 | FEMALE | | | band | |
| 4294-012 | FEMALE | | | band | |
| 4294-013 | FEMALE | | | band | |
| 4294-014 | FEMALE | | | band | |

5' Homology Arm:
Cyp1a2 stop KI gen F CACAAAAGGAACACAAAGGAAAGG
dTAG 5HA R tcagtttggctctctgaccc
Product size: 624 bp targeted only

3' Homology Arm:
dTAG 3HA F attcctcccgggacagaaac
Cyp1a2 stop KI gen R CAGGGTAGGAGGAATCTTAAATCA
Product size: 418 bp targeted only

Internal:
Cyp1a2 FKBP F tgcaggtggaaaccatctcc
Cyp1a2 FKBP R ccagtggcaccataggcata
Product size: 253 bp targeted only

**Figure S2. Evaluation of the effects of Cyp1a2$^{dTAG}$ on liver using H&E staining and ALT&AST assay. a** and **b**, H&E staining of WT and HOM mice. **c**, Plasma ALT from WT and *Cyp1a2*$^{dTAG}$-HOM mice. **d**, Plasma AST from WT and *Cyp1a2*$^{dTAG}$-HOM mice (n = 4). WT, wild-type; HOM, homozygous.

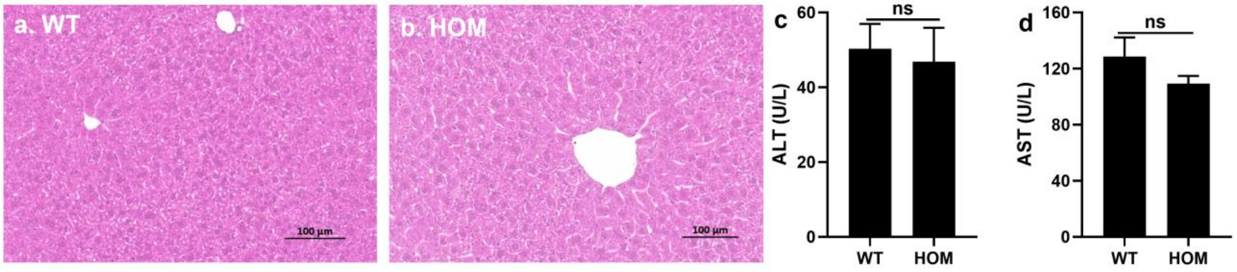

**Figure S3. The expression of CYP1A2 and CYP1A2$^{dTAG}$ in major mouse organs and turnover rate of CYP1A2$^{dTAG}$. a & b,** The expression of CYP1A2 and CYP1A2$^{dTAG}$ in brain, lung, kidney, spleen, testis, and intestines from WT and *Cyp1a2*$^{dTAG}$-HOM mice (n = 2). **c** & **d**, Time course of degradation and turnover rate of CYP1A2$^{dTAG}$ in mouse liver after dTAG-13 treatment (n = 2). **e**, The level of CYP1A2$^{dTAG}$ protein at 24 h after one treatment and two treatments at 0 & 8 hours. (n = 3 for each group). Protein levels in vehicle treated mice were normalized to 100%.

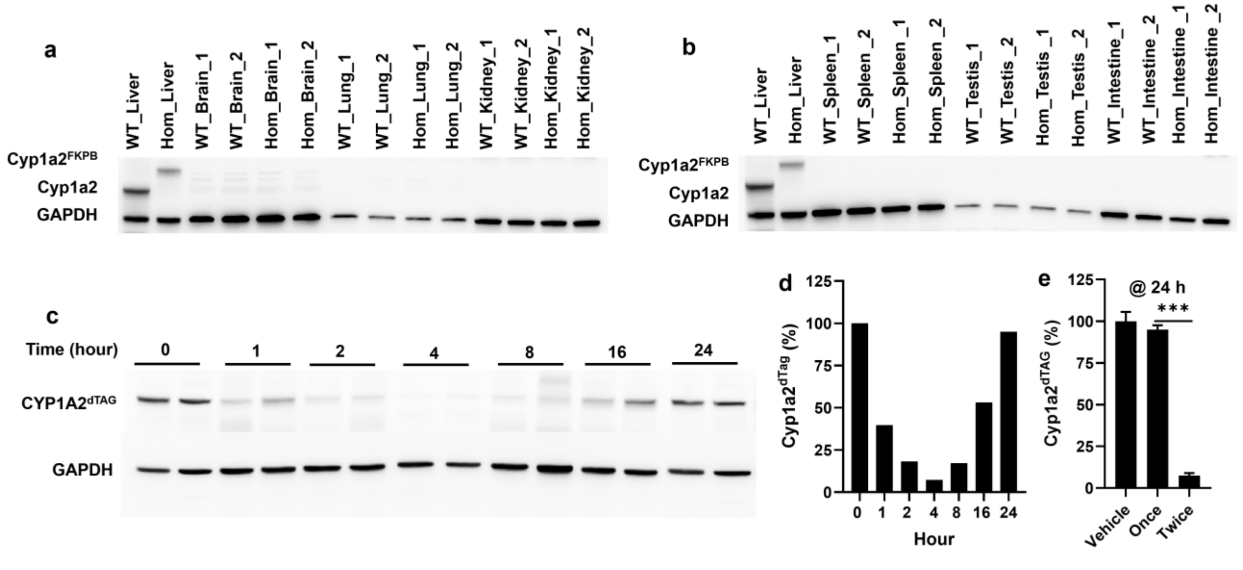

**Figure S4. Relative abundance of mRNA of Cyp1a2, Cyp3a11, Cyp2e1, Ugt1a1, and Abcg2 in WT and *Cyp1a2*$^{dTAG}$ -HOM mice.**

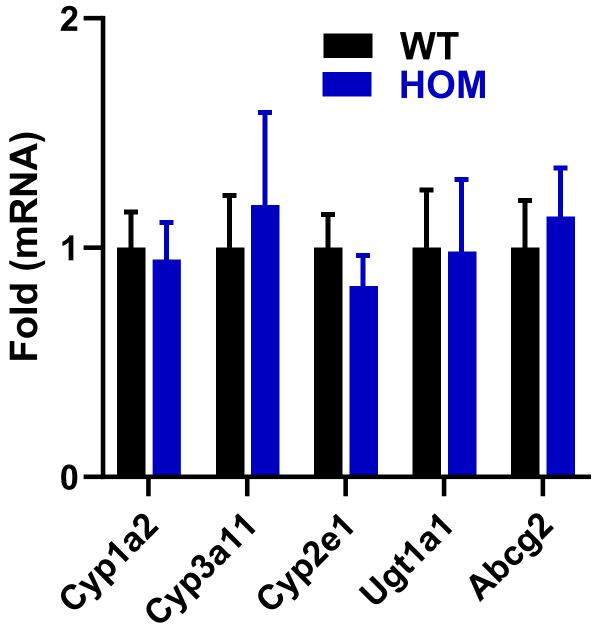

**Figure S5. Proteomics analysis of mouse liver from WT and *Cyp1a2*$^{-/-}$ mice.** (n = 3, females)

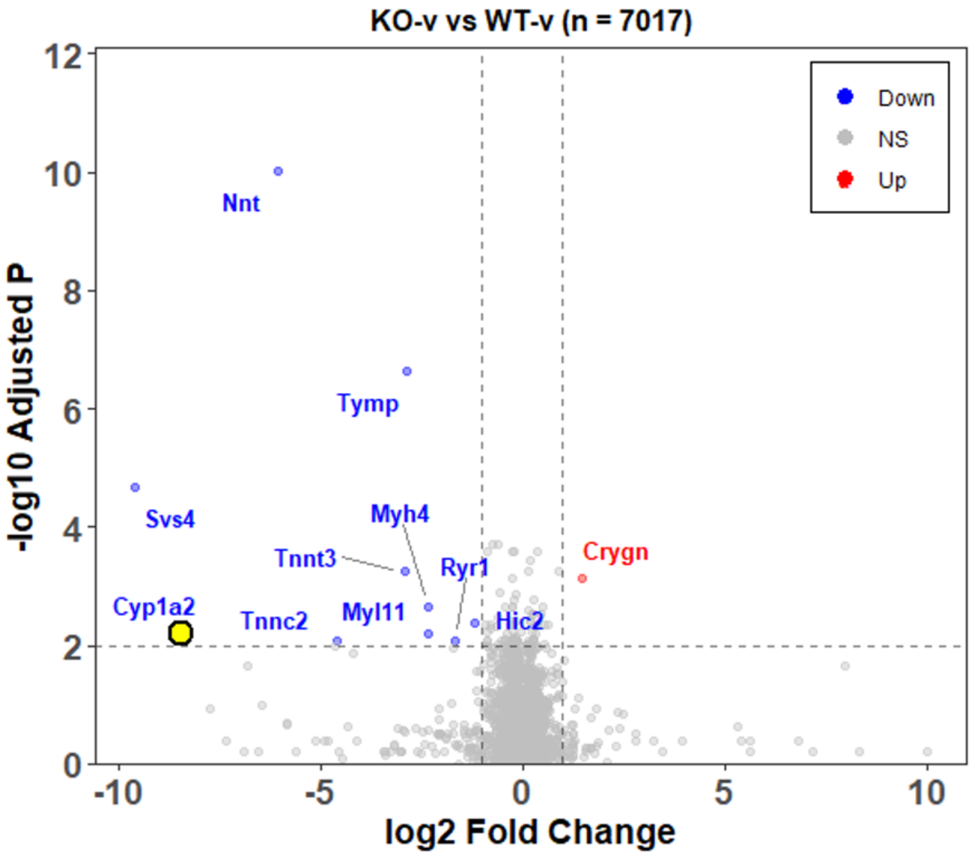

**Figure S6. PK of theobromine and theophylline (Cyp1a2-less involved caffeine metabolites) in *Cyp1a2*$^{\mathrm{dTAG}}$ -HOM mice. a,** Metabolic pathways of caffeine. **b**, PK profile of theobromine in *Cyp1a2*$^{\mathrm{dTAG}}$-HOM mice treated with vehicle or dTAG-13. **c**, PK profile of theophylline in *Cyp1a2*$^{\mathrm{dTAG}}$-HOM mice. The caffeine (10 mg/kg, p.o.) was administered to mice 4 hours post vehicle or dTAG-13 (20 mg/kg, i.p.). After two-week washout and recovery following PK of caffeine in mice treated with vehicle, PK in mice treated with dTAG-13 was performed. (n = 6).

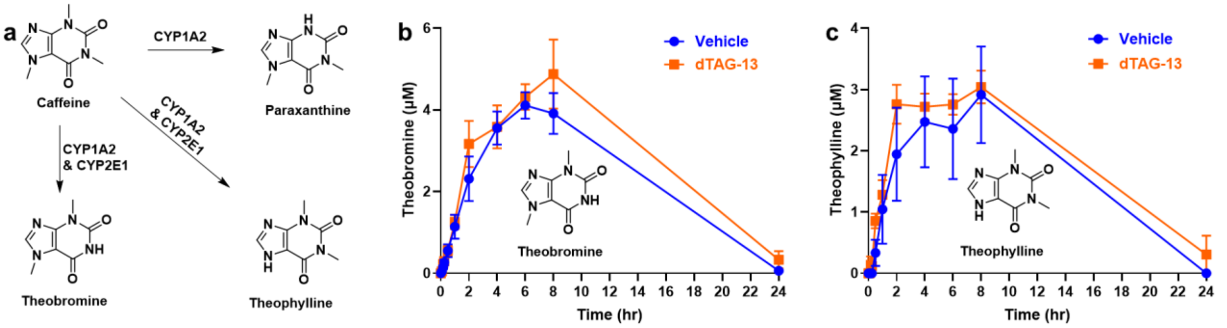

**Figure S7. Design of *Cyp3a11* dTAG knock-in.** The *Cyp3a11* N terminus will be targeted using two gRNAs flanking the translational start site to insert the $FKBP^{F36V}$ sequence and flexible linker. PCR genotyping will be used to identify targeted alleles [primers will give 5' and 3' HA products of 584 and 495 bp (respectively) for the dTAG allele]. The PCR product from the flanking primers will be 629 bp or 965 bp for WT or dTAG, respectively.

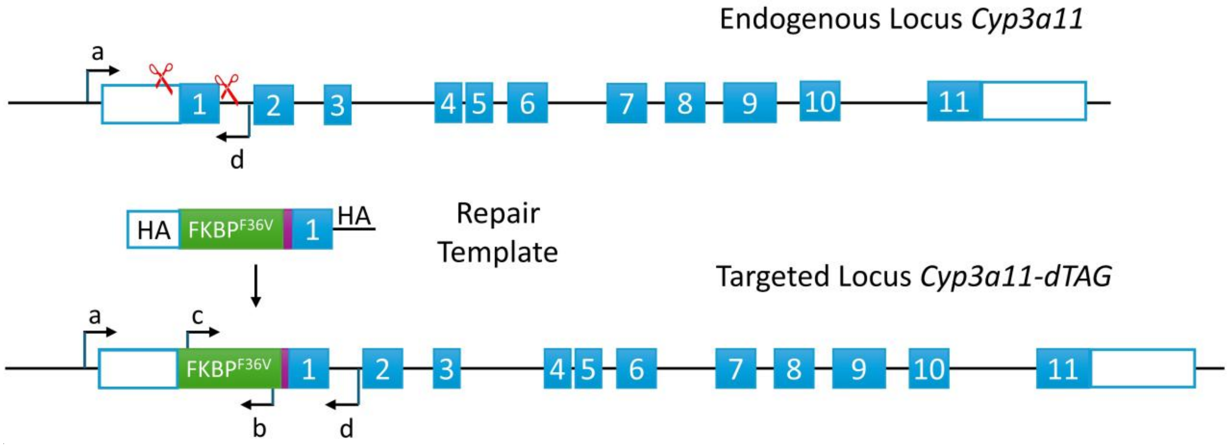
